# Melanin-concentrating hormone receptor antagonism differentially attenuates nicotine experience-dependent locomotor behavior in female and male rats

**DOI:** 10.1101/2023.08.31.555726

**Authors:** Isabel R.K. Kuebler, Youxi Liu, Bárbara S. Bueno Álvarez, Noah M. Huber, Joshua A. Jolton, Raaga Dasari, Ken T. Wakabayashi

**Affiliations:** Neurocircuitry of Motivated Behavior Laboratory, Department of Psychology, University of Nebraska–Lincoln, 1220 T St. Lincoln, Nebraska 68588; Rural Drug Addiction Research Center, University of Nebraska–Lincoln, 660 N 12th St. Lincoln, Nebraska 68588

**Keywords:** sex differences, locomotion, sensitization, tobacco use disorder, non-contingent administration, hypolocomotion

## Abstract

Nicotine is a significant public health concern because it is the primary pharmacological agent in tobacco use disorder. One neural system that has been implicated in the symptoms of several substance use disorders is the melanin-concentrating hormone (MCH) system. MCH regulates various motivated behaviors depending on sex, yet little is known of how this interaction affects experience with drugs of abuse, particularly nicotine. The goal of this study was to determine the effect of MCH receptor antagonism on experience-dependent nicotine-induced locomotion after chronic exposure, particularly on the expression of locomotor sensitization. Adult female and male Wistar rats were given saline then cumulative doses of nicotine (0.1, 0.32, 0.56, and 1.0 mg/kg) intraperitoneally to determine the acute effects of nicotine (day 1). Next, rats were treated with 1.0 mg/kg nicotine for 6 days, given an identical series of cumulative doses (day 8), and then kept in a drug-free state for 6 days. On day 15, rats were pretreated with vehicle or the MCH receptor antagonist GW803430 (10 or 30 mg/kg) before another series of cumulative doses to assess response to chronic nicotine. After vehicle, male rats increased nicotine locomotor activation from day 1 to day 15, and both sexes showed a sensitized response when normalized to saline. The lower dose of GW803430 decreased locomotion compared to vehicle in females, while the higher dose decreased locomotion in males. Both sexes showed nicotine dose-dependent effects of GW803430, strongest at lower doses of nicotine. Controlling for sex-based locomotor differences revealed that females are more sensitive to GW803430. The high dose of GW803430 also decreased saline locomotion in males. Together, the results of our study suggest that MCH is involved in the expression of nicotine locomotor sensitization, and that MCH regulates these nicotine behavioral symptoms differently across sex.

## 1 Introduction

Nicotine, the primary pharmacological agent causing tobacco use disorder (TUD), continues to present substantial public health concerns, despite recent promising declines in nicotine use (Office of the Surgeon General, 2014). For example, smoking nicotine-containing tobacco remains the top cause of preventable death and disease in the United States (Office of the Surgeon General, 2014). Further, 47.1 million Americans, or 19% of adult men and women, used one or more nicotine-containing products including cigarettes, e-cigarettes, cigars, pipes, and smokeless tobacco as of 2020 (Cornelius, 2022). Moreover, the use of e-cigarettes has received increased attention because the recent use of this product among high school students grades 9-12 has reached a disproportionate 14.1% (CDCTobaccoFree, 2022). Quitting the use of these products remains a personal and public health challenge. Particularly, continued smoking is prevalent among people wanting to quit (Leventhal et al., 2022) despite several smoking cessation medications available. Smoking and smoking cessation patterns show wide differences across sex, suggesting neurobiological differences in response to nicotine and pharmacological treatments (Cornelius, 2022; P. H. Smith et al., 2016). Therefore, novel pharmacological approaches may add to current treatments to aid in successful smoking cessation. One way to achieve this goal may be by targeting novel neural systems beyond those historically studied in TUD.

One potential therapeutic target that has been increasingly shown to influence symptoms of substance use disorders like TUD is the melanin-concentrating hormone (MCH) system. Historically, MCH has been studied in the context of energy homeostasis, sleep-wakefulness, feeding, alcohol intake, and psychostimulant reinforcement and neurobehavioral adaptations (Barson & Leibowitz, 2016; Chung et al., 2009; Diniz & Bittencourt, 2017; Duncan et al., 2005, 2006; Georgescu et al., 2005; Rossi et al., 1997; Sun et al., 2013; Verret et al., 2003). Despite being implicated in several use disorders, a systematic examination of MCH and nicotine has not been undertaken. Determining the effects of MCH function on nicotine-induced behavior may further uncover contributions of MCH’s regulation toward TUD-like behavior.

One commonly studied behavioral symptom of nicotine is locomotor activity after acute and chronic nicotine experience. With drugs like cocaine and amphetamine, a key behavioral symptom of repeated drug exposure in rodent preclinical models is a sensitization of the locomotor activating effects of the drug (Robinson & Berridge, 1993). This development is correlated with increased self-administration, faster acquisition of self-administration, motivation to work for, and greater propensity toward reinstatement of a drug (Covington & Miczek, 2001; De Vries et al., 1998; Horger et al., 1992; Lorrain et al., 2000; Mendrek et al., 1998; Piazza et al., 1990; Vezina, 2004). Importantly, locomotor sensitization after drug withdrawal has been associated with increases in drug-induced craving, a key symptom of substance use disorder (Lu et al., 2004; Shaham et al., 2003). Behavior resembling sensitization to psychostimulants has been reported by some groups studying the effects of nicotine, with low locomotion after acute nicotine and increasing levels after repeated exposure (Booze et al., 1999; Ericson et al., 2010; Kanýt et al., 1999; Ksir, 1994). Degrees of change in locomotion over time vary between research groups, which may be due to the wide range of challenge doses of nicotine used in different studies, between 0.1 mg/kg and 4.0 mg/kg in rats (Benwell et al., 1995; Domino, 2001). One commonly used method used to determine a full dose response within a single session, and thereby reducing the number of rats used in an experiment is cumulative dosing (Schechter, 1997; Wenger, 1980). Therefore, in this study, we determined changes in locomotor activity across different doses by using cumulative dosing within a single session, an approach that allows us to determine the behavioral effects of nicotine at multiple doses within a session. An additional benefit to these designs is the ability to incorporate a saline injection before the onset of nicotine dosing. This allowed us to assess changes in baseline locomotor responses associated with the experimental procedure, such as habituation to the open field locomotor chamber.

In addition, varying patterns of nicotine-induced locomotion can become more prevalent once sex as a biological variable is considered. For example, many studies report increased behavioral activation over time in females when compared to males (Booze et al., 1999; Hamilton et al., 2014), but others report similar behavior between sexes (Ericson et al., 2010; Honeycutt et al., 2020). Further, locomotor activation reaches a maximum at moderate doses of nicotine, and high doses of nicotine induces low locomotion for both sexes (Elliott et al., 2004). This may be attributed to the activation of low-affinity nicotinic acetylcholine receptors at high doses of nicotine (Picciotto, 2003; Tizabi et al., 2000). In animal models of passive exposure and self-administered nicotine, females show different patterns of rewarding as well as aversive conditioned responses to nicotine (Torres et al., 2009) and self-administer more nicotine than males (Flores et al., 2019). As well, MCH also impacts sex-specific behaviors and energy regulation depending on sex. For example, MCH regulates feeding more in male than female rats (Santollo & Eckel, 2008; Terrill et al., 2020). Yet, there has been a dearth of reports examining MCH function, its relationships with nicotine-induced activity, and sex as a biological variable thus far. This is surprising because with other drugs of abuse, such as cocaine and amphetamine, other research groups have shown that MCH function has a complex effect on drug-induced locomotor activation (Chung et al., 2009; D. G. Smith et al., 2008; Sun et al., 2013).

As only one of two MCH receptor subtypes, MCH receptor 1 (MCHR1), is expressed in rats (Tan et al., 2002), the objectives of this study were to examine broadly the effects of MCHR1 antagonism on experience-dependent nicotine-induced locomotion in both female and male rats over a range of nicotine doses. Another goal of this study was to compare the locomotor responses of female and male rats in a passive nicotine administration model originally developed exclusively in male rats. The specific objective of this study was to determine the effect of MCHR1 antagonism on the expression of nicotine locomotor sensitization after drug withdrawal, an important correlate of drug-induced craving. We hypothesized that both male and female rats would dose-dependently increase in locomotion in response to nicotine and that MCHR1 antagonism would attenuate this change. Moreover, we predicted that females would increase behavioral activation to a greater extent than males, shown by greater changes in locomotion, and that MCHR1 antagonism would have a stronger effect in females.

## 2 Materials and Methods

### 2.1 Subjects

Data from 40 male and 39 female age-matched Wistar rats (Envigo) weighing 270-290g (male) or 250-270g (female) upon arrival were included in this study. Rats were co-housed in pairs by sex in a climate-controlled animal colony equipped with filtered air cages. Rats had *ad libitum* access to standard chow (Teklad) and water. Rats were maintained on a 12-12 light/dark cycle with lights on 1500. Rats were acclimatized to the animal facility for at least 3 days before the beginning of experimental procedures. Female rats were intact and free-cycling, as we had no *a priori* hypotheses on the effect of ovarian cycles on sensitization. All procedures were approved by the University of Nebraska–Lincoln Animal Care and Use Committee and complied with the Guide for the Care and Use of Laboratory Animals (National Research Council (U.S.) et al., 2011).

### 2.2 Drugs

Drugs included (-)-nicotine (tartrate) (Cayman, Ann Arbor, Michigan, U.S.) dissolved in sterile 0.9% physiological saline and adjusted to a pH of 7.2-7.4 prior to the injection. All doses of nicotine reported in this study reflect the weight of the base. The MCH receptor antagonist 6-(4-Chloro-phenyl)-3-[3-methoxy-4-(2-pyrrolidin-1-yl-ethoxy)-phenyl]-3*H*-thieno[3,2-*d*]pyrimidin-4-one (GW803430) (Axon Medchem, Groningen, Netherlands) was freshly prepared the day of the treatment, and dissolved in a vehicle solution of 1 part ethanol, 1 part emulphor (Solvay, Brussels, Belgium), and 18 parts saline (Feja et al., 2020; Niphakis et al., 2013).

### 2.3 Nicotine-induced behavioral activation

The procedure for assessing nicotine-induced locomotor activation was modified from Liu et al. 2018, and consisted of six phases: habituation, acute, treatment, post-treatment, withdrawal, and post-withdrawal. An illustration of the general experimental timeline is shown in Figure 1.

**Figure 1.**
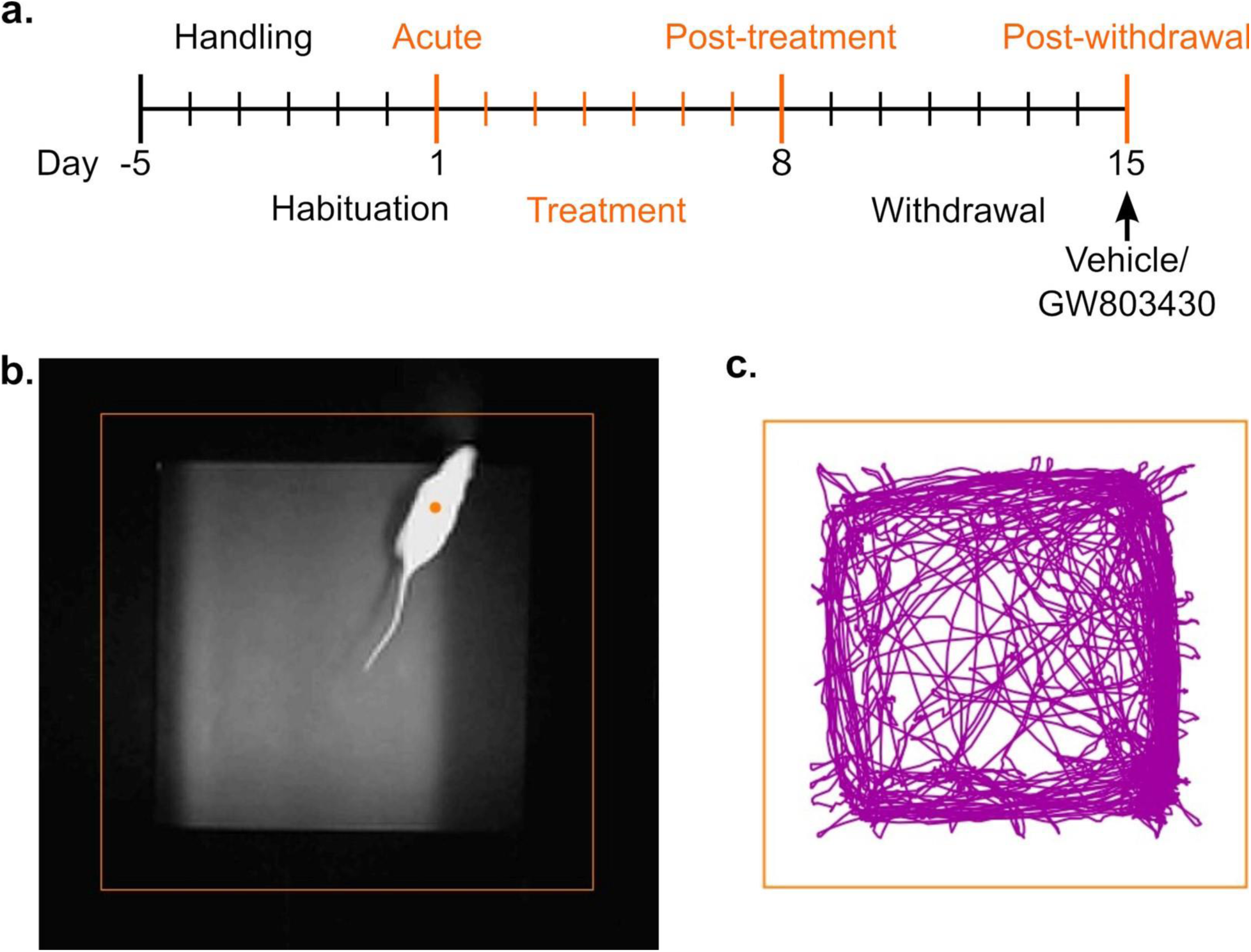
Methods. (a) A timeline of experimental procedures with nicotine treatments shown in orange. Tick marks represent each day, with the first large tick mark indicating arrival. Large orange tick marks indicate cumulative dosing sessions, with a pretreatment of vehicle or GW803430 30 minutes before the last session. (b) Example image of an open field chamber with computer derived orange tracking borders and orange computer derived tracking dot in the center of the rat’s body. (c) Representative tracking plot showing the rat’s locomotor path in purple over the entire 100-minute dose response session.

#### 2.3.1 Habituation phase

After 3 days of handling, rats were habituated to black acrylic open field locomotor chambers (59.5 cm L x 59.5 cm W x 40.6 cm H; Central Plasticworks) 1 hr per day for two days prior to the beginning of experimental procedures. Open fields were equipped with video cameras (Genius WideCam F100) for video tracking.

#### 2.3.2 Acute phase

On day 1 of the experiment, a within session dose-response curve was determined to the acute effects of nicotine during a 100-min session. Rats were first injected with saline (1 mL/kg) intraperitoneally (i.p.), and then placed in the open field chamber. Subsequently, every 20 minutes, rats were injected at cumulative doses of 0.1, 0.32, 0.56, and 1.0 mg/kg nicotine (Liu et al., 2018) and video recorded to assess acute locomotor response to nicotine. The specific doses used to achieve these cumulative doses were 0.1, 0.22, 0.24, and 0.44 mg/kg, respectively.

#### 2.3.3 Treatment phase

Starting the next day, on days 2-7 of the experiment, rats were treated with 1.0 mg/kg (i.p.) of nicotine once per day. During these sessions, locomotion was video recorded for 1 hour.

#### 2.3.4 Post-treatment phase

On day 8, a nicotine dose-effect curve identical to the acute session (Day 1) was conducted to assess the post-treatment effects of nicotine.

#### 2.3.5 Withdrawal phase

On days 9-14, rats were kept in a drug-free state for 6 days in their home cages. During this phase all rats were regularly handled every 2-3 days.

#### 2.3.6 Post-withdrawal phase

Rats were assigned treatment groups by their overall locomotion from day 8 of the experiment (post-treatment phase) so that all of the groups were balanced. On day 15 of the experiment, rats were pretreated with either vehicle or GW803430 (10 mg/kg or 30 mg/kg, i.p.) 30 minutes before the session and placed in the open field chamber. Then, a nicotine dose effect-curve was redetermined to assess changes in the locomotor response to nicotine.

### 2.4 Data validation and analysis

Locomotion during the experiment was assessed by video tracking (ANY-maze version 7.0). All raw video recordings and the subsequent automated tracking was carefully examined by at least 2 experimenters. Videos were screened for experimental artifacts such as accidental mis-tracking of clothing, hands, and the fidelity of the tracking itself. To aid in removing tracking artifacts, locomotion data was split into bins of 20 seconds. Bins containing visible video artifacts or inaccurate tracking were removed from subsequent analyses. Potential outliers for each treatment group of the post-withdrawal dose response locomotion were flagged if a subject’s data fell outside of the 1.5 interquartile range (IQR) from the median for that dose. Identified subjects were then carefully inspected for their experimental history, to determine any possible experimental errors and assess the overall reliability of the data for inclusion. The data was considered validated once two raters reached a consensus to include these data for further analysis.

Locomotion was measured as total distance (m) traveled for 15 minutes after each dose, starting 5 minutes after the injection to omit initial locomotion in response to the injection (Li et al., 2013). All results are presented as mean ± standard error of the mean (SEM).

Statistical analysis was completed using GraphPad Prism version 9/10 and RStudio version 4.2.2. At first, locomotion (as distance traveled) between sexes at each session was determined by two-way repeated measures Mixed-Effects Analysis with nicotine dose matched within subjects, and sex as the between-subjects factor. During our analyses, we determined that saline locomotion during the acute, post-treatment and post-withdrawal session was significantly different (Fig. 3a, two-way repeated measures Mixed-Effects Analysis, *F*_2,110_=13.08, *p*<0.0001), suggesting that rats were habituating to the experimental procedure over time. To account for this, we also normalized the locomotion at each dose of nicotine as percent change from saline locomotion on that day before follow-on two-way repeated measures Mixed-Effects Analysis with nicotine dose matched within subjects, and sex as the between-subjects factor.

Locomotion in response to nicotine during the post-withdrawal phase (day 15) was determined within each sex via two-way repeated measures Mixed-Effects Analysis with saline and each dose of nicotine matched within subjects, and treatment as the between-subjects factor. As females and males treated with vehicle varied in locomotor activation after nicotine experience, each treatment group was normalized to vehicle as percent change from the vehicle mean at that dose of nicotine or saline. Normalized nicotine locomotion data was analyzed via three-way repeated measures Mixed-Effects Analysis with each dose of nicotine matched within subjects and sex and treatment as between-subject factors. Treatment and sex were then subsequently analyzed within each dose of nicotine via two-way analysis of variance (ANOVA).

Finally, the effect of MCH receptor antagonism on saline locomotion within and between each sex during the post-withdrawal phase was directly compared via two-way ANOVA.

Post-hoc multiple comparisons for each analysis were adjusted for comparing between sexes or treatment groups using the Holm-Šídák method or Dunnett’s method when comparing against control values. For Mixed-Effects Analyses, we used a compound symmetry covariance matrix, fitted using Restricted Maximum Likelihood (REML). For clarity, only significant effects and interactions are statistically detailed. The level of significance was set to α=0.05.

## 3 Results

### 3.1 Response to nicotine over different experimental sessions differs by sex

We found that sex had a markedly different and nuanced influence on nicotine-induced locomotion depending on the experimental session. We first compared locomotor activation as distance traveled between female and male rats during each session. During the acute session (day 1), females had higher locomotion than males overall, especially at low doses of nicotine, and males had similar locomotion at each nicotine dose so that there was a fixed effect of nicotine dose (Fig. 2a, two-way repeated measures Mixed-Effects Analysis, *F*_3,231_=31.39, *p*<0.0001) and sex (*F*_1,77_=25.45, *p*<0.0001) and a nicotine dose x sex interaction (*F*_3,231_=19.93, *p*<0.0001). We then compared locomotor activity between female and male rats after repeated nicotine treatment. Females had overall similar levels of locomotion compared to males but higher locomotion at the lowest dose, (Fig. 2b, fixed effect of dose of nicotine *F*_3,227_=43.17, *p*<0.0001), an interaction of nicotine dose x sex (*F*_3,227_=9.195, *p*<0.0001), but no effect of sex. Next, we compared locomotion between sexes after the drug-free post-withdrawal period. Females demonstrated higher overall locomotion than males particularly at the lowest dose so that there was a fixed effect of nicotine dose (Fig. 2c, *F*_3,99_=42.53, *p*<0.0001), a fixed effect of sex (*F*_1,33_=7.973, *p*=0.0080), and an interaction of nicotine dose x sex (*F*_3,99_=4.194, *p*=0.0077). Taken together, nicotine-induced locomotion generally was higher at lower doses of nicotine and decreased at the higher doses, and female rats had greater locomotion than male rats at low doses of nicotine over all sessions.

**Figure 2.**
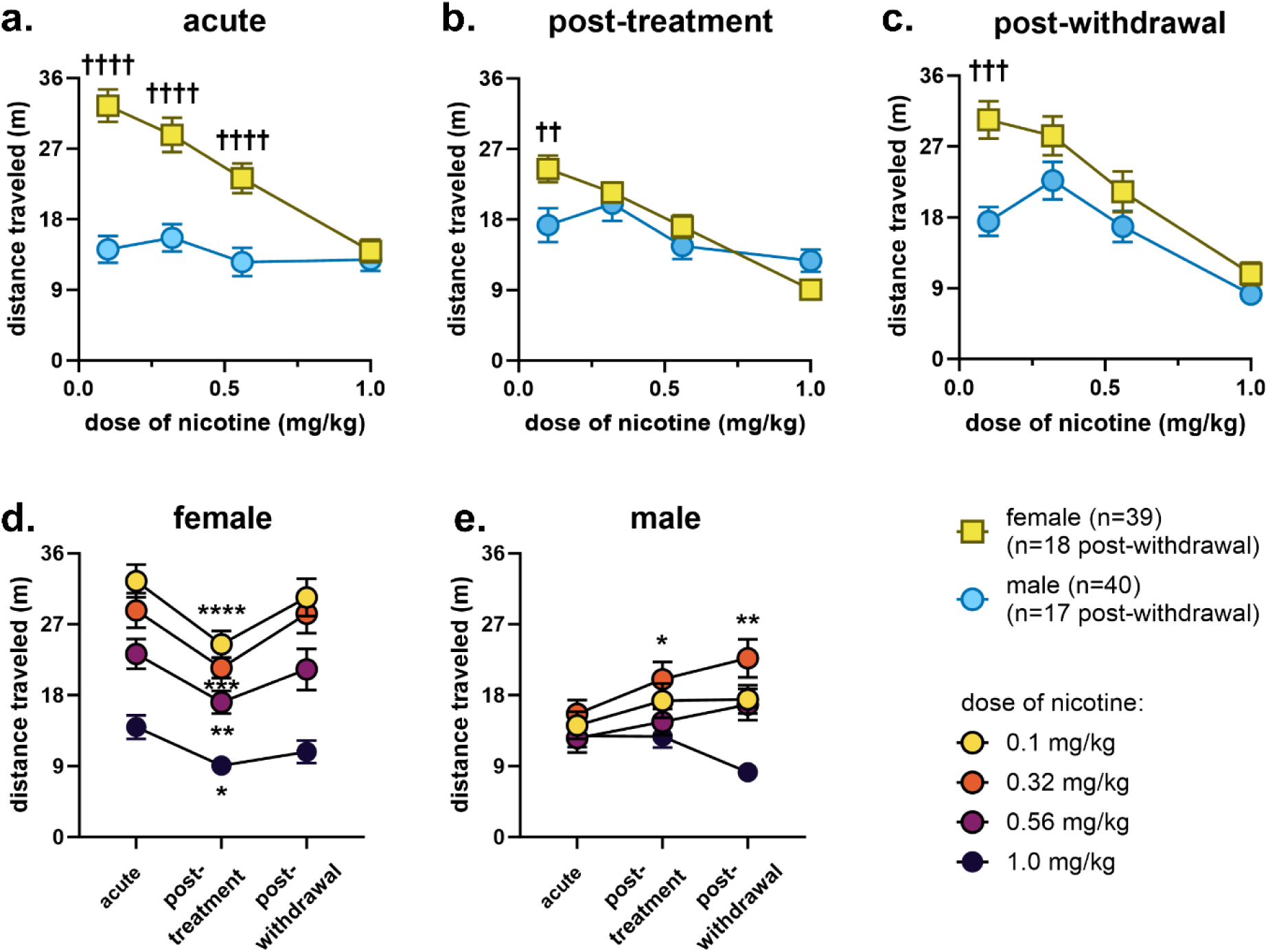
Locomotor response (as distance traveled) to nicotine over different treatment sessions differs strongly by sex. All data are shown as mean ± SEM. (a) Females generally decreased in locomotion as nicotine dose increased in the acute session, while males show similar locomotion at each dose of nicotine; locomotion was greater in females at the three lowest doses of nicotine. (b) Females show greater locomotion than males at the lowest dose of nicotine post-treatment. (c) Female rats again show greater locomotion than males at the lowest dose of nicotine post-withdrawal (d) Female rats show a dip in locomotion at all doses of nicotine between the acute and post-treatment sessions and a return to acute levels post-withdrawal. (e) Males show increases in locomotion compared to the acute session at a moderately low dose of nicotine. †† p < 0.01, ††† p < 0.001, †††† p < 0.0001 by Holm-Šídák’s multiple comparisons test. * p < 0.05, ** p < 0.01, *** p < 0.001, **** p < 0.0001 compared to acute by Dunnett’s multiple comparison test.

We then compared the locomotor reactions of each sex between the acute (day 1) session and the subsequent sessions. For female rats there was a fixed effect of session (Fig. 2d, two-way repeated measures Mixed-Effects Analysis, *F*_2,220_=24.61, *p*<0.0001) and a fixed effect of nicotine dose (*F*_3,152_=34.04, *p*<0.0001), but no interaction. There was significantly less locomotion at the post-treatment (day 8) session compared to the acute (day 1) session at every dose of nicotine, but no differences between acute (day 1) and post-withdrawal (day 15). In male rats, there was a fixed effect of session (Fig. 2e *F*_2,216_=4.711, *p*=0.0099) and nicotine dose (*F*_3,156_=6.392, *p*=0.0004) and a session x nicotine dose interaction (*F*_6,216_=2.662, *p*=0.016), so that males showed an increase from the acute (day 1) session for both sessions at 0.32 mg/kg nicotine. Overall, female rats showed consistent decreases from the acute (day 1) session to the post-treatment (day 8); males showed increases between the acute (day 1) session and both later sessions. These male specific increases were stronger at lower doses of nicotine.

Since our saline response during each session indicated a habituation to the experimental session (Fig. 3a, Fixed effect of session, two-way repeated measures Mixed-Effects Analysis, *F*_2,110_=13.08, *p*<0.0001, no effect of sex or a sex x session interaction), we normalized the locomotion at each dose of nicotine as percent change from saline locomotion on that day. During the acute session, as a percent change from saline, females had higher locomotion than males between 0.1-0.56 mg/kg nicotine so there was a fixed effect of nicotine dose (Fig. 3b, two-way repeated measures Mixed-Effects Analysis, *F*_3,231_=30.29, *p*<0.0001) and sex (*F*_1,77_=20.17, *p*<0.0001) and a sex x nicotine dose interaction (*F*_3,231_=18.69, *p*<0.0001). During the post-treatment session, females had higher normalized locomotion than males at the lowest dose of nicotine (Fig. 3c, fixed effect of nicotine dose *F*_3,227_=44.62, *p*<0.0001; sex x nicotine dose interaction *F*_3,227_=10.81, *p*< 0.0001). During the post-withdrawal session, females had higher locomotion accounting for saline locomotion than males overall, and this difference was greatest at the lowest nicotine dose (Fig. 3d, fixed effect of nicotine dose *F*_3,99_=42.70, *p*<0.0001; sex *F*_1,33_=5.844, *p*=0.0213; sex x nicotine dose interaction *F*_3,99_=3.852, *p*=0.0118). Interestingly, this analysis shows that nicotine across all cumulative doses during the acute session induced a suppression below that of the saline control in male rats.

**Figure 3.**
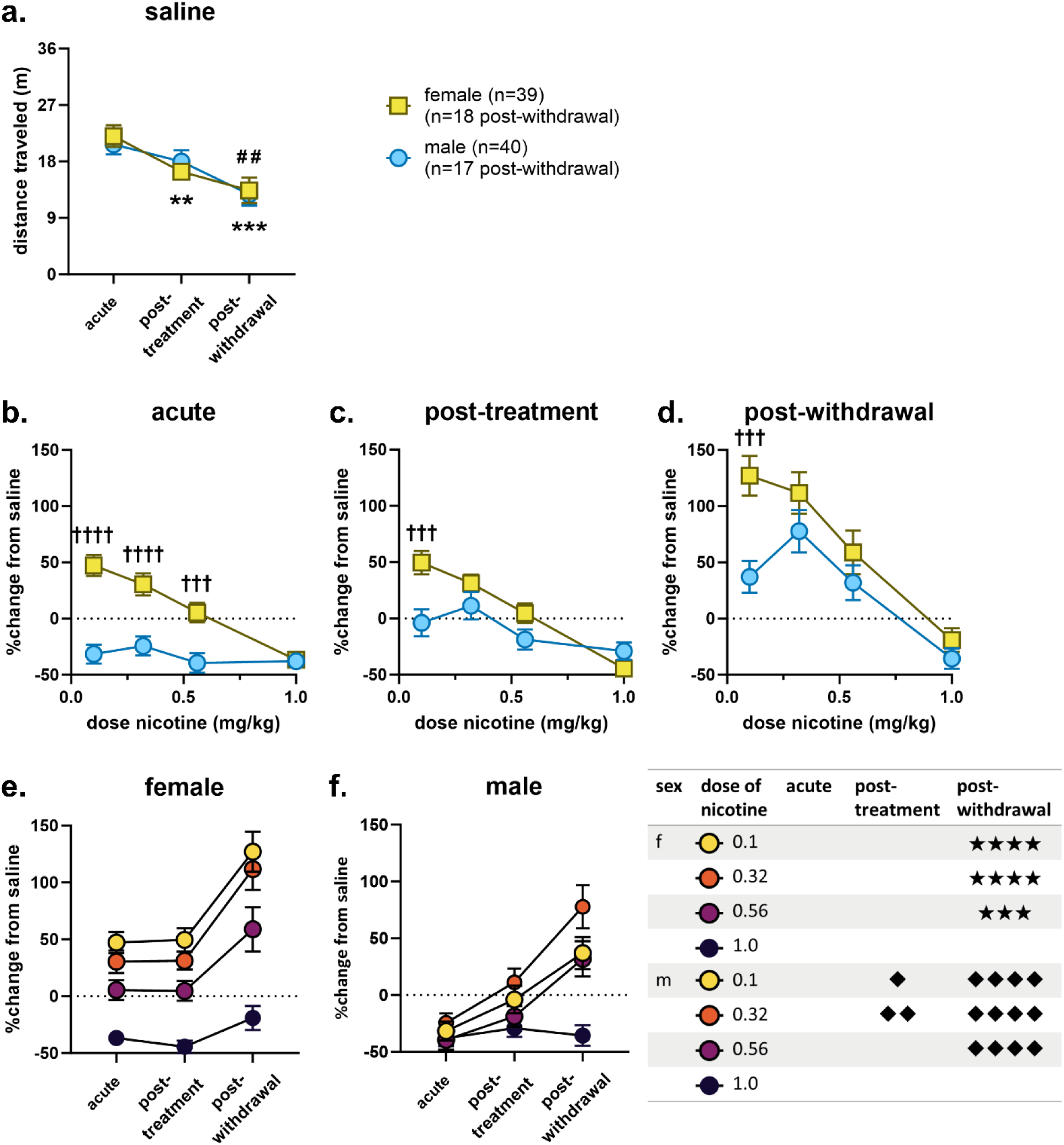
Locomotor response to nicotine normalized to saline locomotion over different treatment sessions increases across session differently by sex. All data are shown as mean ± SEM. (a) Both males and females decreased in saline locomotion across each session. (b) Females generally decreased in saline-normalized locomotion as nicotine dose increased during the acute session, while males show similar locomotion at each dose of nicotine which was lower than the locomotor response to saline for that session; saline-normalized locomotion in females was greater than males at the three lowest doses of nicotine. (c) Females show greater saline-normalized locomotion than males at the lowest dose of nicotine post-treatment. (d) Female rats again show greater saline-normalized locomotion than males at the lowest dose of nicotine post-withdrawal. (e) Between treatment sessions, female rats show similar saline-normalized locomotion during the acute and post-treatment sessions and show a significant increase in saline-locomotion between the acute and post-withdrawal sessions. (f) Males show increases in saline-normalized locomotion between the acute and post-treatment sessions at the two lowest doses of nicotine and an increase in normalized locomotion between the acute and post-withdrawal sessions for the three lowest doses of nicotine. ††† p < 0.001, †††† p < 0.0001 by Holm-Šídák’s multiple comparisons test. ** p < 0.01, *** p < 0.001, **** p< 0.0001; υ p < 0.05, υυ p < 0.01, υυυυ p< 0.0001; ★★★ p < 0.001, ★★★★ p< 0.0001; male saline: ## p < 0.01 compared to acute by Dunnett’s multiple comparison test.

Comparing the saline-normalized locomotion between each session (Fig. 3e, f), there was a consistent increase in nicotine-induced locomotion between treatments for both female and male rats. For female rats, the three lowest doses of nicotine induced significantly greater locomotion between the acute and post-withdrawal sessions (Fig. 3e, two-way repeated measures Mixed-Effects Analysis, fixed effect of dose, *F*_3,114_=109.0, *p*<0.0001; session *F*_2,76_=23.71, *p*<0.0001; nicotine dose x session interaction *F*_6,144_=3.484, *p*=0.0030). In females, no significant changes were seen between the acute and post-treatment sessions. In male rats, all but the highest dose of nicotine induced greater saline-normalized locomotion between the acute and post-withdrawal sessions (Fig. 3f, fixed effect of nicotine dose *F*_3,117_=35.91, *p*<0.0001; session *F*_2,78_=17.80, *p*<0.0001; nicotine dose x session interaction *F*_6,138_=8.756, *p*<0.0001). In male rats, a sensitized locomotor response was detected at the two lowest doses of nicotine between the acute and post-treatment sessions, while a sensitized response was more consistently observed across more doses during the post-withdrawal session. Therefore, once taking into account changes in saline locomotion, a sensitized locomotor response to nicotine was most consistently observed in both sexes during the post-withdrawal session.

We next determined the effects of pretreating rats with the MCHR1 antagonist GW803430 on saline and nicotine locomotion during the post-withdrawal (day 15) session. During this session, there were significant fixed effects of nicotine in both females and males, (Fig. 4, two-way repeated measures Mixed-Effects Analysis, fixed effect of nicotine dose, females (a): *F*_4,144_=38.69; males (b): *F*_4,148_=18.37, both p<0.0001) and MCHR1 antagonist treatment, (females (a): *F*_2,36_=3.505, *p*=0.0407; males (b): *F*_2,37_=4.573, *p*=0.0168). There was a significant nicotine dose x GW803430 interaction in males (Fig. 4b, *F*_8,148_ = 2.264. *p*=0.0259) but not females. Therefore, we found that MCHR1 antagonism decreases the locomotor response to nicotine during the post-withdrawal session in both females and males.

**Figure 4.**
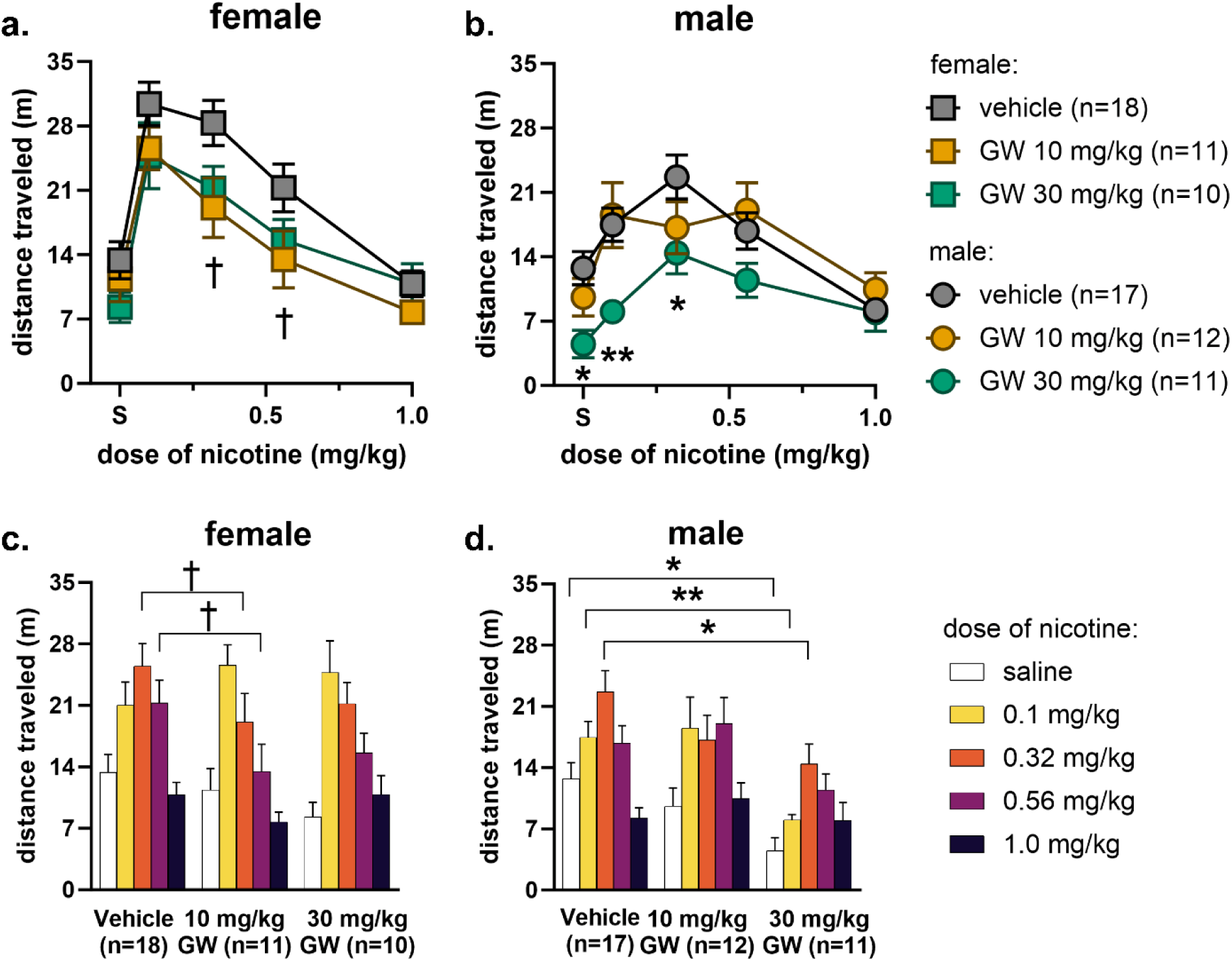
Locomotor stimulation (in distance traveled) after withdrawal from nicotine is attenuated by GW803430. All data are shown as mean ± SEM. For figures a and b, data are shown across saline and different cumulative doses of nicotine. (a) Females decreased in locomotion during the post-withdrawal session (day 15) for each dose of GW803430 compared to vehicle. The lower dose of GW803430 resulted in significantly lower locomotion at 0.32 mg/kg and 0.56 mg/kg of nicotine compared to vehicle. (b) Males decreased in locomotion with 30 mg/kg GW803430 compared to vehicle. 30 mg/kg GW803430 had the greatest effect at the two lowest doses of nicotine, 0.1 and 0.32 mg/kg, compared to vehicle. Males showed an inverted U-shaped dose-response curve after each treatment, while females did not for doses of nicotine. For clarity, the same data from a and b are shown in figure c and d across different doses of GW803430. † p < 0.05, GW803430 10 mg/kg, * p < 0.05, ** p < 0.01, GW803430 30 mg/kg compared to vehicle by Dunnett’s multiple comparisons test.

Post-hoc analyses revealed that for females, there was a significant decrease from vehicle pretreatment control at the lower dose of GW803430 at 0.32 mg/kg and 0.56 mg/kg nicotine (Fig. 4a). In males (Fig. 4b), the 30 mg/kg GW803430 pretreatment dose had significant effects compared to vehicle at saline and low doses of nicotine. When viewing the same data grouped by pretreatment doses, it becomes clearer that while MCHR1 antagonism results in similar locomotor trends after both doses of GW803430 in females (Fig. 4c), in males, robust effects of the MCHR1 antagonist are limited to only the higher dose (Fig. 4d).

### 3.2 Change in locomotion from vehicle during post-withdrawal session depends on sex and dose of GW803430

Males pretreated with vehicle during the post-withdrawal (day 15) session displayed an overall lower locomotor response than similarly pretreated females, especially at low doses of nicotine (Fig. 2c). Therefore, to better compare the effects of MCHR1 antagonism between sexes during this session, we normalized the nicotine-induced locomotor response after each dose of GW803430 as percent change from vehicle. There was a fixed effect of nicotine dose (Three-way Mixed-Effects Analysis, *F*_3,120_=3.279, *p*=0.0234), but no significant effects of sex or pretreatment. There was a significant sex x pretreatment interaction (*F*_1,40_=6.153, *p*=0.0174). Follow-up two-way ANOVAs within each dose of nicotine revealed a main effect of pretreatment (Fig. 5a, two-way ANOVA, *F*_1,40_=5.737, *p*=0.0214) and a sex x pretreatment interaction (Fig. 5a, *F*_1,40_=4.819, *p*=0.0340) at 0.1 mg/kg of nicotine. Post-hoc comparisons within each pretreatment dose were not significant across sex.

**Figure 5.**
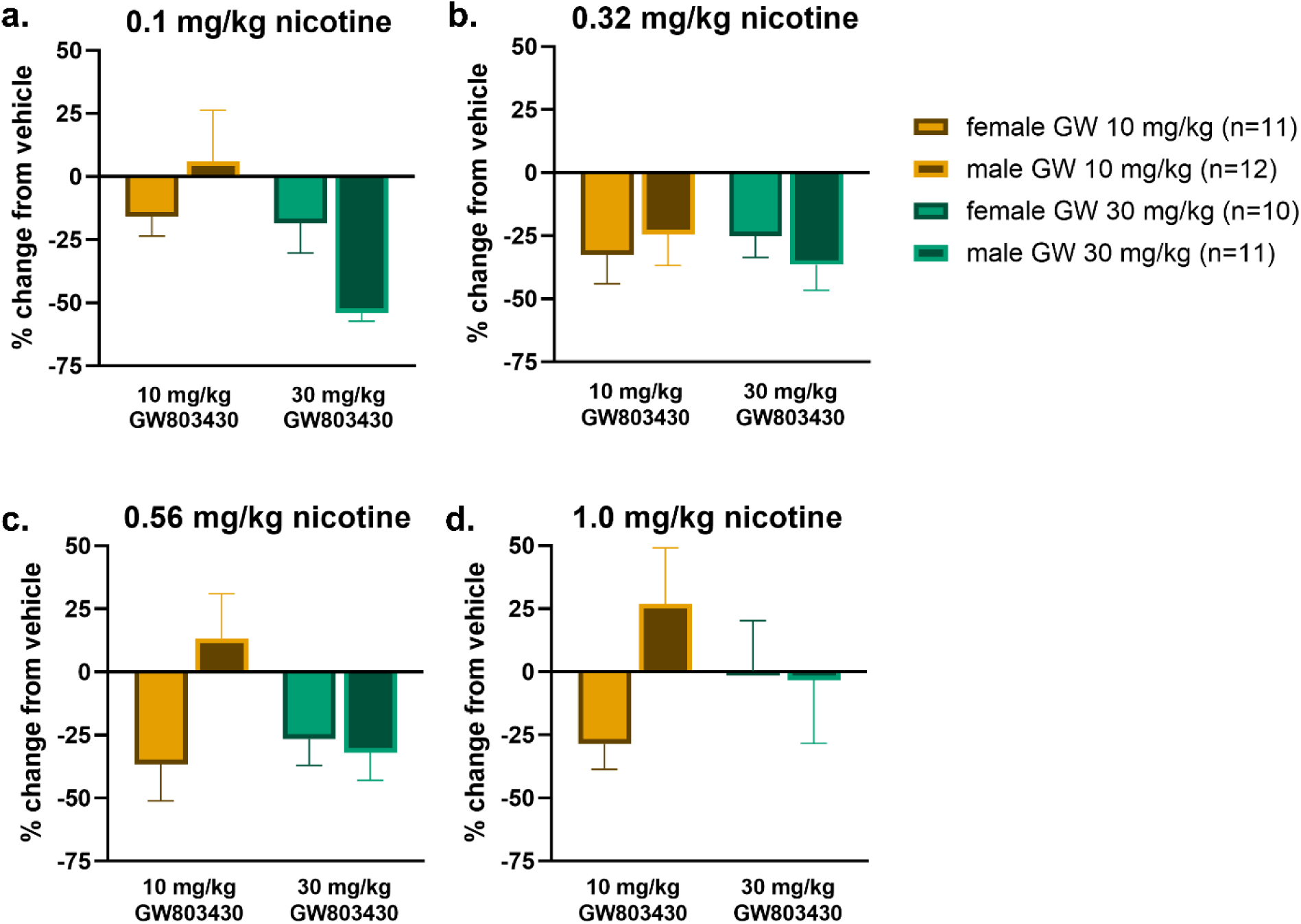
Locomotion changes from vehicle during the post-withdrawal session vary by sex at each dose of nicotine at the same treatment. All data are shown as mean ± SEM. (a) At 0.1 mg/kg of nicotine, when normalized to percent change from vehicle control, 30 mg/kg GW803430 decreased locomotion overall compared to 10 mg/kg GW803430. Males decreased more between 10 and 30 mg/kg GW803430 than females, which show similar responses at each dose. (b-d) Neither dose of GW803430 shows significantly different locomotion between females and males at 0.32, 0.56, or 1.0 mg/kg of nicotine.

To better encapsulate the effect of MCHR1 antagonism on nicotine-induced locomotion during the post-withdrawal (day 15) session, we integrated locomotion for each GW803430 pretreatment dose over all cumulative doses of nicotine as area under the curve (AUC; i.e. [locomotion * nicotine dose]) and compared each subject’s treatment effects as a difference from the vehicle mean. MCHR1 antagonism decreased overall nicotine-induced locomotion at both doses of GW803430 for females (Fig. 6, One sample *t* test, *p*=0.0096, 10 mg/kg, and *p*=0.0094, 30 mg/kg) and only the higher dose of GW803430 in males (Fig. 6, One sample *t* test, *p*=0.0029). Again, females show an effect at a lower dose of GW803430 than males.

**Figure 6.**
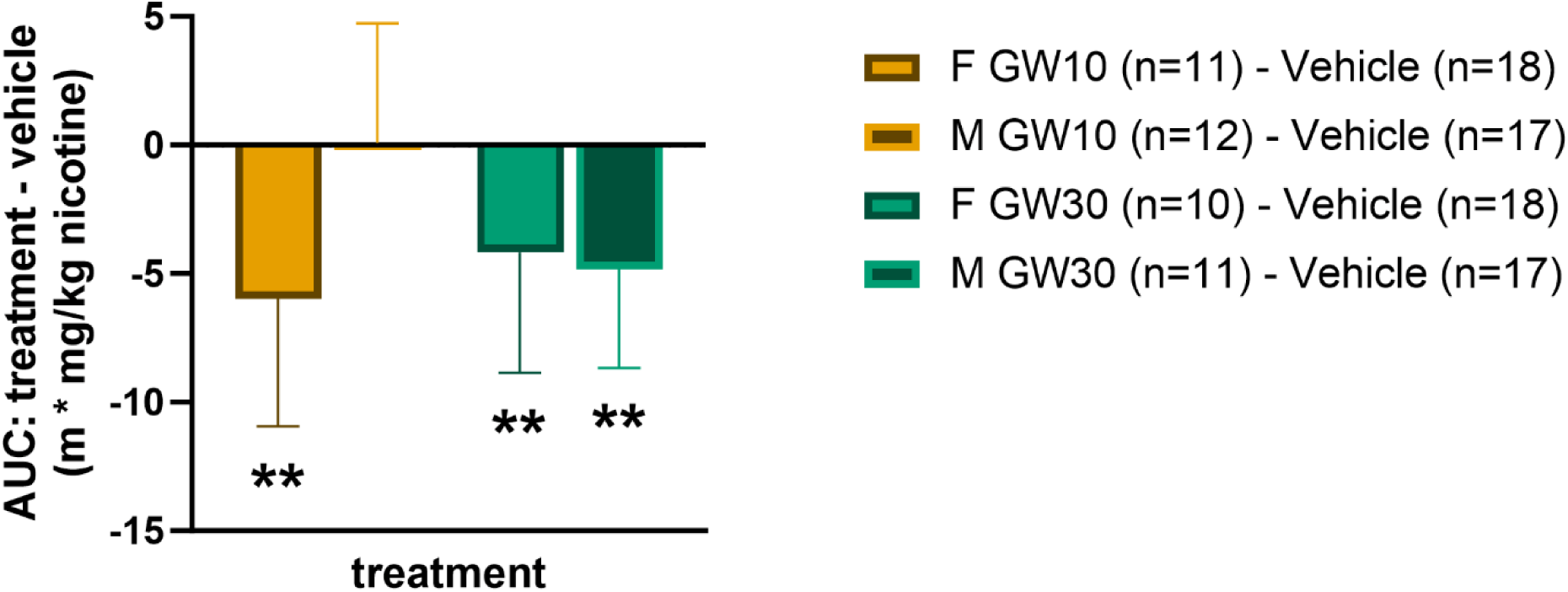
Integration of all doses of nicotine as Area Under the Curve show decreases from vehicle after both doses of GW803430 in females and only after the higher dose of GW803430 in males. Bars and whiskers show mean ± SEM. ** p<0.01 compared to 0 by one-sample t test.

### 3.3 MCHR1 antagonism decreases overall locomotion after saline in males

In our experimental design, the within-subjects dose response procedure also included a saline control injection prior to cumulative doses of nicotine. Somewhat surprisingly, when we analyzed the impact of MCHR1 antagonism on saline locomotion within the omnibus test, there was a post-hoc effect in males (Fig. 4). Out of an abundance of caution, we further assessed this effect by itself. We found that MCHR1 antagonist treatment had an overall effect on locomotion across sex for a main effect of treatment (Fig. 7, two-way ANOVA, *F*_2,73_=4.875, *p*=0.0103). There was no main effect of sex or sex x treatment interaction. Post-hoc comparisons revealed that males, but not females, showed a statistically significant dose-dependent decrease in baseline locomotion. Of note, saline locomotion after GW803430 vehicle pretreatment was similar between females and males.

**Figure 7.**
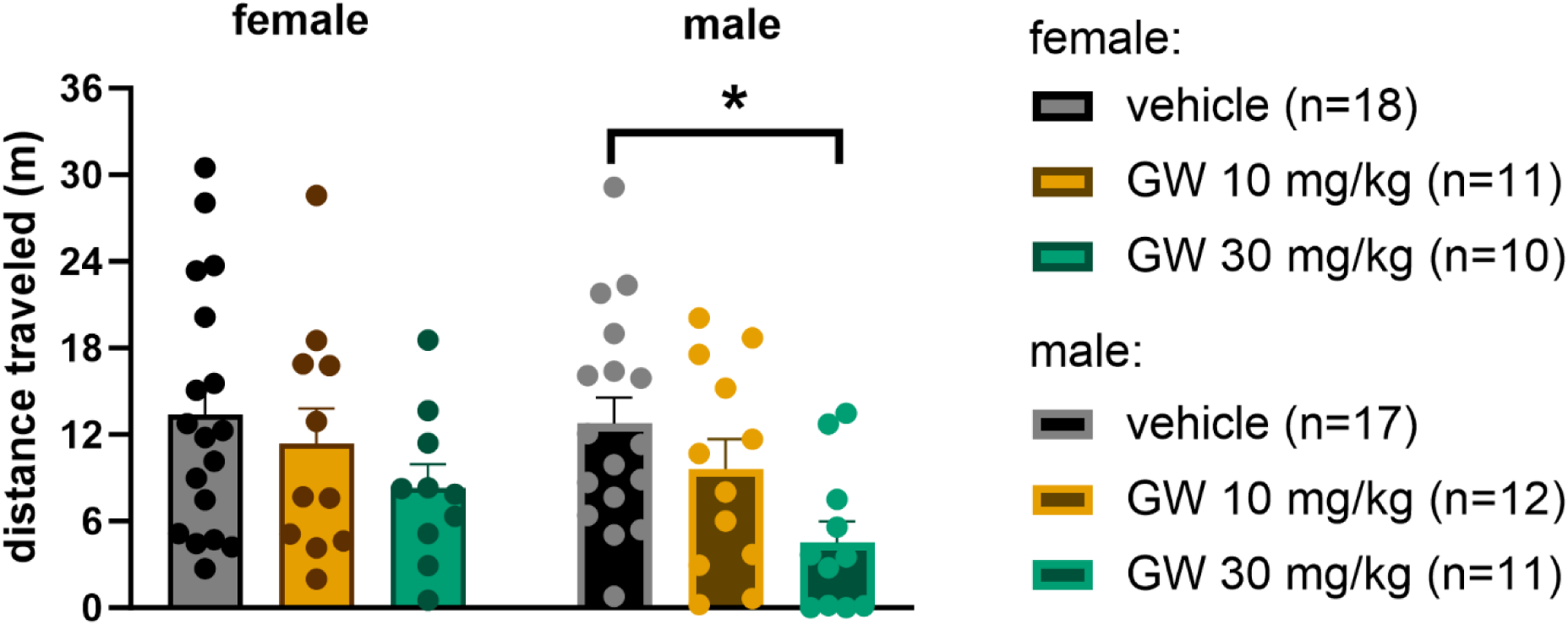
Saline locomotion after pretreatment of GW803430 is decreased in males but not in females. Bars and whiskers show mean ± SEM, points show individual data. Saline locomotion was not significantly different between each treatment group in females. Saline locomotion decreased at the higher dose of GW803430 in males. Only the higher dose of GW803430 in males showed a change from vehicle. * p < 0.05 by Dunnett’s multiple comparisons test.

## 4 Discussion

The objective of this study was to determine the effect of MCHR1 antagonism on the expression of nicotine locomotor sensitization after drug withdrawal in female and male rats. When analyzed directly as distance traveled, males showed a typical increase in locomotor stimulation over time while females showed a more complex pattern of locomotor responses. During our experiment, our saline locomotor response during our cumulative dosing suggested that habituation to the experimental procedure was a factor. Once we accounted for this influence, both females and males showed increases in saline-normalized nicotine-induced locomotion over time. Our novel findings were that systemic pretreatment with the MCHR1 antagonist GW803430 during the post-withdrawal session (day 15) dose dependently decreased a sensitized nicotine-induced locomotor response in both male and female rats, and that these responses also depended on sex. Specifically, females at the lower dose of GW803430 decreased in nicotine-induced locomotion, while in males this effect was limited to the higher dose of GW803430. We also found an effect of GW803430 on saline locomotion at the higher dose in males.

### 4.1 Locomotor responses to nicotine over time differ between sexes

We initially hypothesized that both female and male rats would show an increase in locomotor responses to nicotine between the acute (day 1) and post-withdrawal (day 15) sessions, as typically seen in males given this schedule of nicotine experience (Liu et al., 2018). In our study, male rats displayed a dose-dependent heightened locomotor response across the three sessions. This effect fits the canonical development of locomotor sensitization, an increase in the locomotor activating effects of the same dose of drug (Stewart & Badiani, 1993). This symptom is hypothesized to reflect neuroadaptive behavioral changes associated with disrupted motivation for nicotine and other TUD-like behaviors after repeated exposure (Covington & Miczek, 2001; Piazza et al., 1990; Robinson & Berridge, 1993). Surprisingly, the pattern differed in females. When analyzed directly as distance traveled, female rats showed less locomotion immediately after daily nicotine treatment and then only reached levels of locomotion similar to the initial acute response to nicotine after the drug-free period. Therefore, a strict pharmacological interpretation of this behavior pattern in females would be that they did not develop locomotor sensitization to nicotine. However, one benefit to the cumulative dosing experimental design employed here is the incorporation of a within-subjects saline control injection, particularly since nicotine manipulations may potentiate the rat’s habituation to an open field (Schildein et al., 2002). Our results indicated that open field habituation was a factor in our experiment. Once we normalized our nicotine locomotion to the saline response within the same session, we found a strong, dose-, session-, and sex-dependent sensitized locomotor response to nicotine.

One factor that may affect female rats differently from males is the treatment dose of nicotine in this procedure, which was previously developed and optimized to induce nicotine sensitization exclusively in male rats (Liu et al., 2018). As we used a relatively high dose, this likely impacts the timing and magnitude of changes in locomotor stimulation in females more than more moderate doses of nicotine. It should be noted that, high doses of psychostimulants can result in stereotypy which is also associated with low levels of locomotion (Flagel et al., 2008; Ksir, 1994); qualitative observations during the nicotine treatment portion of the current study generally noted hypolocomotion rather than behavior more indicative of stereotypy, like repetitive movements of the head. In our study, male rats exhibited an inverted U-shaped curve post-withdrawal (day 15), and female rats displayed only the descending limb of the dose-response curve. This suggests that nicotine induced hypolocomotion at higher doses for all subjects during the last session. Another factor that may explain the differences in locomotion is that female rats are more sensitive to the effects of some drugs than male rats over a range of physiological and behavioral measures (Becker & Koob, 2016; Donny et al., 2000; Kuebler et al., 2022). For instance, female rats show higher levels of arterial nicotine than males given the same dose (Harrod et al., 2007), suggesting greater vulnerability to the drug because of pharmacophysiological mechanisms. Certainly, more work will be required to determine the effects of nicotine in female rats and the elements specific to female-specific behavioral responses.

### 4.2 MCHR1 antagonism attenuates nicotine-induced locomotion by dose of nicotine and dose of GW803430

In our study, we tested the effect of the MCHR1 antagonist GW803430 on locomotor responses to cumulative doses of nicotine within a single session while rats were in nicotine withdrawal. Taking the total effect of all doses of nicotine together (Fig. 6), only the low dose of GW803430 decreased overall locomotion compared to the vehicle control in female rats, while only the high dose of the drug had an effect in male rats. This overall effect suggests that MCHR1 antagonism fundamentally impacts the locomotor effects of nicotine and attenuates the expression of sensitization at a range of doses of nicotine. Further, it suggests that normal MCH function may be critical for the expression of nicotine sensitization. Looking within each sex, only the low dose of GW803430 significantly reduced nicotine-induced locomotion at two moderate doses of nicotine from vehicle in females (Fig. 4a). Male rats showed a nicotine dose-dependent effect only at the higher dose of GW803430 (Fig. 4b), with statistically significant effects at saline and 0.1 mg/kg and 0.32 mg/kg of nicotine. Therefore, only the high dose of GW803430 showed significant decreases in nicotine-induced locomotion at low to moderately low doses of nicotine in males, while in females, the low dose of GW803430 showed significant decreases only at moderately low doses of nicotine. Overall, we found that systemic MCHR1 antagonism attenuates nicotine-induced expression of locomotor sensitization in both male and female rats, and that these effects differ depending on both the dose of nicotine and the antagonist.

### 4.3 MCHR1 antagonism decreases nicotine-induced locomotion differently across sex

In order to directly compare the effect of MCHR1 antagonism across sexes, we normalized for differences in baseline locomotor responses to nicotine on the post-withdrawal day (day 15, Fig. 2c). Once accounting for each sex’s locomotor response to nicotine itself, males and females had different responses to the high and low doses of GW803430 (Fig. 5). Females showed similar magnitude decreases in nicotine locomotor effects after both low and high MCHR1 antagonist doses compared to males. In general, only the higher dose of GW803430 was effective in males in reducing nicotine-induced locomotion over a range of nicotine doses. It should be noted that at the moderately low dose of nicotine (0.32 mg/kg), both doses of GW803430 were equally effective in reducing locomotion in both males and females. Together, controlling for locomotor differences in response to nicotine itself across sex revealed that female rats are more sensitive to the effects of GW803430 than males. These results show that sex is an important variable that impacts how effective GW803430 is in reducing dose-dependent effects of nicotine-induced locomotion.

### 4.4 Sex-specific nicotine-independent locomotor responses to MCHR1 antagonism

In females, decreases in locomotor response with GW803430 only occurred after nicotine and not after saline (Fig. 7), suggesting that in females, GW803430’s effects are limited to reducing the effects of nicotine and not motor responses in general. On the other hand, we found that GW803430 decreased the saline locomotion in males compared to vehicle at the higher dose of GW803430 (Fig. 7), suggesting that in males, GW803430 also has dose-dependent effects on nicotine-free locomotion. While the baseline effect of GW803430 might suggest that MCHR1 antagonism in males may function via a different mechanism than in females, this could be a sex-specific side effect of MCHR1 antagonism in general or GW803430 in particular. We were surprised by the reduction of saline locomotion after MCHR1 antagonism because other MCH manipulations not related to drugs of abuse, such as male diet-induced obese mice given chronic GW803430 (Zhang et al., 2014) or female mice with a partial genetic knockout of MCHR1 (Chee et al., 2019) have resulted in increases in locomotion. It should be noted that while our within-subjects cumulative dosing experimental design imparts significant benefits compared to a discrete between-subjects design, our experiment lacks a completely untreated saline control (Schechter, 1997; Wenger, 1980) While historically such an untreated saline control has been reported (Gui-Hua et al., 1992; Schechter, 1997), this approach is uncommon presently (Li et al., 2013; Liu et al., 2018; Thorn et al., 2014; Wu et al., 2020; Xue et al., 2019). Moreover, considering that nicotine experience alone may interact with open-field habituation (Schildein et al., 2002), having an untreated saline control may introduce an unintended confound. Admittedly, why MCHR1 antagonism reduces saline locomotion preferentially in male but not female nicotine-sensitized rats will need to be explored further.

### 4.5 Neurobehavioral interactions of MCH function and effects of drugs of abuse

The results of our study suggest that MCHR1 antagonism attenuates the expression of nicotine sensitization in both sexes, in good agreement with studies examining the interaction of MCH and other drugs of abuse. Our findings are also relevant because the expression of nicotine sensitization during drug withdrawal is viewed as a behavioral correlate of the incubation of drug craving, a key diagnostic symptom of TUD and other forms of substance use disorders (American Psychiatric Association, 2022; Lu et al., 2004; Shaham et al., 2003). Anatomically, MCH neurons project to both the nucleus accumbens and the ventral tegmental area (Diniz et al., 2019), indicating that MCH modulates motivated behaviors regulated by the mesolimbic system, the primary circuit regulating reward-related behavior. There is increasing evidence for this hypothesis. First, MCH activity generally heightens alcohol seeking and intake (Cippitelli et al., 2010; Duncan et al., 2005; Morganstern et al., 2010). MCH has a more complex effect on psychostimulants, with MCH function associated with either increased (Chung et al., 2009) or attenuated sensitization (D. G. Smith et al., 2008; Sun et al., 2013). Historically, nicotine’s neurobiological impact on the mesolimbic system has been considered a fundamental mechanism associated with the symptoms of TUD. However, the neurobiological mechanism underlying the impact of nicotine administration on MCH function remains unclear. On the one hand, nicotinic acetylcholine receptor antagonism in the hypothalamus results in biochemical downregulation of lateral hypothalamic MCH neurons (Garcia et al., 2015). In contrast, Jo and colleagues found that MCH neurons in the lateral hypothalamus receive indirect cholinergic input. The authors predict that a nicotine challenge would inhibit MCH activity (Jo et al., 2005). Taken together, more information is needed to determine the precise mechanism of nicotine’s effects on MCH circuits contributing to nicotine sensitization.

### 4.6 Sex as a biological variable impacts the effects of MCH function and nicotine experience

Our study shows that MCH regulates symptoms associated with nicotine across sex, contributing to the growing understanding that sex differences are integral to MCH function. For example, others have found that MCH regulates feeding via MCHR1 binding in the nucleus accumbens in male rats but does not regulate feeding in female rats (Terrill et al., 2020). We provide initial evidence that MCH shows sex differences with a drug of abuse. Specifically, our study demonstrates that female rats are more sensitive to the effects of MCHR1 antagonism than male rats. Our study further emphasizes the importance of sex differences in the response to nicotine alone and its regulation by the MCH system.

### 4.7 Future directions

In addition to the effects of MCHR1 antagonism on nicotine-induced locomotion, we found differences from male locomotor stimulation patterns in female rats using a high treatment dose of nicotine. Determining the effects of MCHR1 antagonism during nicotine treatment sessions, particularly considering a range of treatment doses and treatment times will also be valuable in uncovering interactions of MCH function and nicotine effects. Finally, the impact of MCH manipulations on nicotine self-administration and associated models of relapse would provide further generalizable information as well.

## 5 Conclusions

Our study demonstrated that MCHR1 antagonism dose-dependently attenuates nicotine-induced locomotion in male and female rats. In addition, our findings suggest that MCH regulates the expression of nicotine locomotor sensitization, particularly when accounting for habituation to the experimental procedure. Controlling for sex-specific locomotion levels, both sexes showed a nicotine dose-dependent effect of MCH receptor antagonism, generally responding most strongly at low doses of nicotine. Female rats were more sensitive to the antagonist than males, suggesting an important sex difference. In all, these results implicate the MCH system in symptoms central to TUD. Therefore, targeting MCH function may be a promising therapeutic avenue for TUD in humans.

## Acknowledgments

The authors thank M. Matuszeski, K. Sextro, and G. Zimmerman for assistance with experimental procedures and data validation, and J. Kuebler and Drs. R. Bevins and M. Li for helpful comments on an early draft of the manuscript.

